# FLAgellum Member 8 modulates extravascular trypanosome distribution in the mammalian host

**DOI:** 10.1101/2021.01.08.425862

**Authors:** Estefanía Calvo Alvarez, Christelle Travaillé, Aline Crouzols, Brice Rotureau

## Abstract

The African trypanosome flagellum is an essential organelle in multiple aspects of the parasites’ development. Here, we investigated the role of a flagellar protein termed FLAgellar Member 8 (FLAM8) that is specifically distributed along the entire flagellum in trypanosomes stages of the vertebrate host. Analyses of knockdown and knockout trypanosomes demonstrated that FLAM8 is not essential *in vitro* for survival, growth, motility and slender to stumpy differentiation. Functional investigations in experimental infections showed that *FLAM8*-deprived trypanosomes are able to establish and maintain the infection in the blood circulation, and to differentiate into insect transmissible forms. However, quantitative bioluminescence imaging revealed that *FLAM8*-null parasites exhibit an impaired dissemination in the extravascular compartment, that is partially restored by the addition of a single rescue copy of *FLAM8*. Interestingly, among all dissected organs scrutinized individually, only the skin of mice infected with *FLAM8*-deprived parasites showed a significant reduction in extravascular trypanosome population as compared to mice infected with parental controls. To our knowledge, FLAM8 is the first example of a flagellar protein that modulates *T. brucei* parasite distribution in the host tissues, contributing to the maintenance of extravascular parasite populations in mammalian anatomical niches, especially in the skin.

**Take away:**

- FLAM8 is dispensable *in vitro* for survival, growth, motility and differentiation of *T. brucei*.
- FLAM8 depletion does not affect parasitemia and bloodstream form differentiation *in vivo*.
- FLAM8 modulates the extravascular dissemination of trypanosomes in the mammalian host, especially in the skin.

## Introduction

*Trypanosoma brucei* is an extracellular parasite responsible for African trypanosomiases in sub-Saharan Africa, a group of neglected tropical diseases including sleeping sickness in humans and nagana in cattle. African trypanosomes are blood and tissue-dwelling protists transmitted by the bite of the blood-feeding tsetse fly (*Glossina* genus). In the mammalian host, parasites have to face different micro-environments, including deadly challenges by multiple types of host immune responses and the variable availability of carbon sources. This requires major morphological and metabolic adaptations, driven by the activation of specific gene expression programs, that are critical for life-cycle progression (MacGregor, Szoor, Savill, & Matthews, 2012; Ooi & Bastin, 2013; Smith, Bringaud, Nolan, & Figueiredo, 2017). Recently, the importance of extravascular tropism for *T. brucei* has been re-discovered and emphasized in animal models: in addition to the brain, parasites actually occupy most mammalian tissues, especially the skin and the adipose tissues (Caljon et al., 2016; Capewell et al., 2016; Trindade et al., 2016). However, when, where and how trypanosomes adapt to survive within these extravascular environments is not understood yet.

The trypanosome flagellum is an essential organelle anchored along the surface of the cell body and present in all stages of its development (Rotureau, Subota, & Bastin, 2011). It is essential for parasite viability (Broadhead et al., 2006), cell division and morphogenesis (Kohl, Robinson, & Bastin, 2003), attachment to the tsetse salivary glands (Tetley & Vickerman, 1985) and motility (Langousis & Hill, 2014). In the insect host, the flagellum remains at the forefront of the cell and is likely to be involved in sensory and signaling functions required for host-parasite interactions (Roditi, Schumann, & Naguleswaran, 2016; Rotureau, Morales, Bastin, & Spath, 2009). In the mammalian host, flagellar motility was shown to be critical for establishment and maintenance of bloodstream infection (Shimogawa et al., 2018). Nevertheless, the overall contributions of the trypanosome flagellum to parasite tropism and spatiotemporal dissemination dynamics in the mammalian host are important aspects that still remained to be explored more in depth.

Our proteomic analysis of intact flagella purified from the insect stage of the parasite identified a group of flagellar membrane and matrix proteins with unique patterns and dynamics (Subota et al., 2014). Amongst them, one large protein (3,075 amino acids) termed FLAgellar Member 8 (FLAM8) is present only at the distal tip of the flagellum in the insect procyclic form (Bertiaux, Morga, Blisnick, Rotureau, & Bastin, 2018; Fort, Bonnefoy, Kohl, & Bastin, 2016; Subota et al., 2014). Interestingly, it is redistributed along the entire length of the flagellum in mammalian bloodstream forms (BSF), including in stumpy transmissible stages (Calvo-Alvarez *et al*., in revision), which may imply a specific function for FLAM8 in the vertebrate host. Therefore, we hypothesized that the stage-specific redistribution of FLAM8 could be involved in host-parasite interactions, possibly affecting parasite homing and dynamics in host tissues. Here, we investigated the possible roles of FLAM8 *in vitro* in terms of survival, proliferation, motility and differentiation. In addition, experimental infections in mice monitored by bioluminescence imaging allowed us to decipher the involvement of FLAM8 in parasite dissemination and spreading in both the intra- and extravascular compartments of the mammalian host. Potential consequences for parasite transmission along with cellular mechanisms of tissue tropism are further discussed.

## Results

### *FLAM8* RNAi silencing does not affect parasite survival *in vitro* and *in vivo*

The differential distribution of FLAM8 in the flagellum of the different trypanosome stages (Calvo-Alvarez *et al.*, in revision) raises the question of its specific functions during the parasite cycle. In order to investigate the potential role(s) of FLAM8 in the mammalian host, BSF were first engineered for inducible RNAi knockdown of *FLAM8* in a monomorphic strain expressing a mNeonGreen-tagged version of FLAM8. In order to monitor their behavior in mouse by whole-body imaging approaches, these *FLAM8::mNG FLAM8^RNAi^* mutants were subsequently transformed to overexpress a chimeric triple reporter protein (Calvo-Alvarez, Cren-Travaillé, Crouzols, & Rotureau, 2018). Upon RNAi induction with tetracycline for 72 h, *FLAM8* expression was reduced by 60% at the mRNA level (Fig. 1A) and became undetectable at the protein level by immunofluorescence (Fig. 1B). Parasite growth was monitored over 6 days upon induction of RNAi and no impact on proliferation was observed (Fig. 1C), which indicates that FLAM8 is not essential for survival of BSF parasites in cell culture conditions.

**Figure 1.**
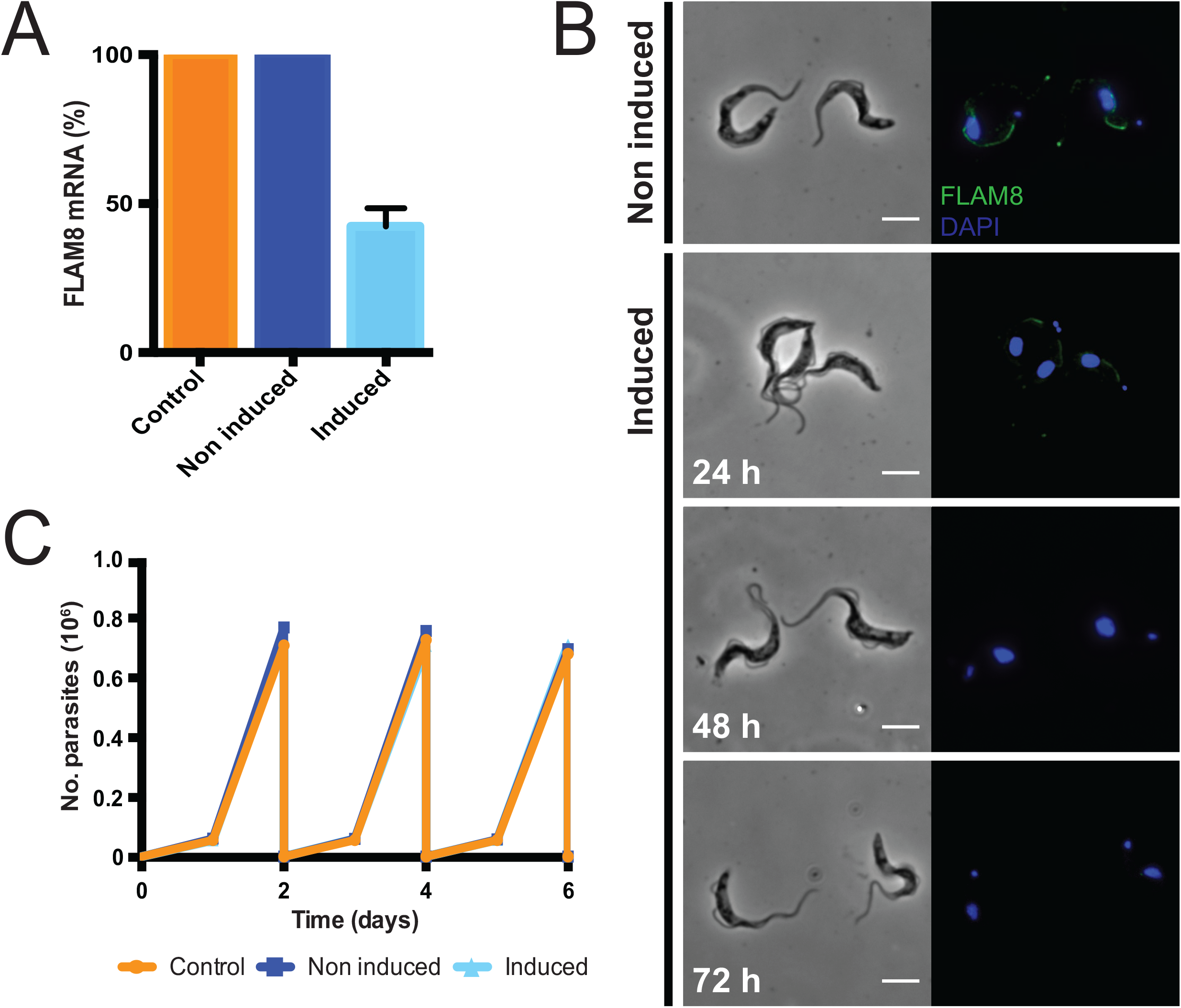
Characterization of the *FLAM8^RNAi^* strains *in vitro*. **A)** Expression of *FLAM8* mRNA assessed by RT-PCR by the comparative ΔΔC_T_ method in control, non-induced and induced *FLAM8^RNAi^* parasites (72 h). **B)** Immunofluorescence pictures of non-induced (upper panels) and induced (bottom panels) *FLAM8^RNAi^ FLAM8::mNG* BSF during 72 h. Methanol-fixed trypanosomes were stained with an anti-mNG antibody (green) and DAPI for DNA content (blue). The scale bars represent 5 μm. **C)** Growth curves of control, non-induced and induced *FLAM8^RNAi^* BSF parasites. All cell lines received 1 μg tetracycline during 6 days. Control parasites are Lister 427 “Single Marker” BSF parasites that do not bear the pZJM-FLAM8 plasmid for RNAi silencing. Results represent the mean (± standard deviation, SD) of three independent experiments.

Then, the linear correlation between the emitted bioluminescence and the number of parasites was assessed in an IVIS Spectrum imager prior to *in vivo* challenge (Fig. S1). In order to get insights into the function of FLAM8 in the mammalian host, groups of male BALB/c mice were infected by the intraperitoneal route with 10^5^ BSF parasites of the parental, non-induced and induced cell lines (Fig. 2). *In vivo* RNAi silencing of *FLAM8* was maintained in mice by the addition of doxycycline in sugared drinking water 48 h prior infection and until the end of the experiment. The course of the infection was monitored daily by i) quantifying the parasitemia, and ii) acquiring the bioluminescent signal emitted by the parasites in the entire organism with an IVIS Spectrum imager. The number of parasites in the extravascular compartment at a given timepoint can be extrapolated by subtracting the known number of trypanosomes in the vascular system (parasitemia x blood volume, according to body weight) from the total number of parasites in the organism (total bioluminescence). No differences were detected neither in the establishment of the infection and the subsequent variations in the number of intravascular parasites (Fig. 2A), nor in the number of parasites occupying extravascular tissues (Fig. 2B) and the animal survival (not shown). Similar population profiles were observed over the course of the infection for both IV and EV parasites in each of the three independent groups of infected mice (Fig. 2, C-E). Finally, in the extravascular compartment, no significant differences were detected in terms of parasite dissemination over the entire animal body in any of the groups tested (Fig. 2, F-G).

**Figure 2.**
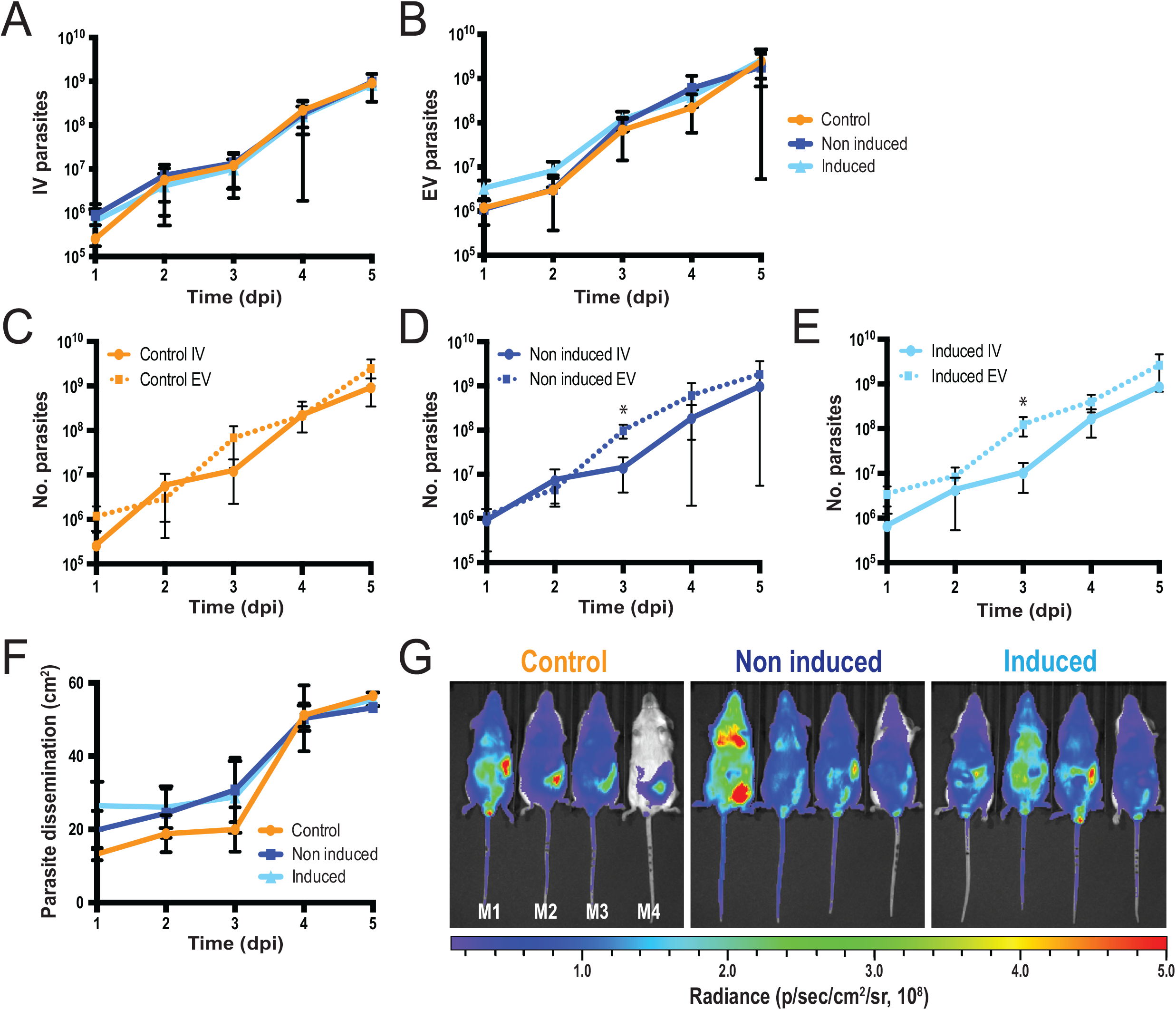
Functional investigations on *FLAM8^RNAi^* parasites *in vivo* in the mammalian host. Groups of 4 BALB/c mice were injected intraperitoneally with either control, non-induced or induced *FLAM8^RNAi^* BSF trypanosomes. One PBS-injected BALB/c animal was used as negative control. **A)** Number of parasites in the blood (intravascular, IV) of infected BALB/c mice during the course of the infection (5 days). **B)** Number of parasites in the extravascular compartment (extravascular, EV) of the same mice. **C, D, E)** Comparison of the number of trypanosomes present in the blood circulation (IV, continuous line) and in extravascular tissues (EV, dotted line) during the entire experimental infection. Significant differences are indicated with * (p<0.05). **F)** Dissemination of control, non-induced and induced *FLAM8^RNAi^* parasites, measured over the entire animal body (in cm^2^) through the total bioluminescent surface, during the entire infection course. Results represent means ± standard deviation (SD). **G)** Representative normalized *in vivo* images of the bioluminescence radiance signal (in photons / second / cm^2^ / steradian) emitted from BALB/c mice infected with control, non-induced and induced *FLAM8^RNAi^* parasites 4 days post-infection (non-infected technical control mice were negative for bioluminescence, not shown).

### *FLAM8* knockout affects trypanosome spreading in the mammalian host

Considering that i) *FLAM8* RNAi silencing efficiency was only partial (40% *FLAM8* mRNA left after 72 h of induction), ii) the efficacy of doxycycline-induced *FLAM8* repression could have been even lower *in vivo*, and that iii) the parental strain used for this first strategy was monomorphic (i.e. unable to differentiate into tsetse adapted stumpy stages), we reasoned that a gene knockout approach in a pleomorphic strain would be more appropriate to evaluate the potential role(s) of FLAM8 during the mammalian host infection. Therefore, a Δ*FLAM8* knockout cell line was generated in pleomorphic BSF trypanosomes by homologous recombination. The replacement of both *FLAM8* alleles by distinct resistance cassettes was verified by whole-genome sequencing and PCRs (Fig. S2, A-B). The trypanosome cell lines generated were further transformed to express the chimeric triple reporter construct as previously described (Calvo-Alvarez et al., 2018), and the linear correlation between the bioluminescence signal and the total number of parasites was analyzed for all strains (Fig. S3). In Δ*FLAM8* knockout parasites where a rescue copy of one *FLAM8* allele was added back into its endogenous locus, the FLAM8 distribution was restored along the entire flagellum, as assessed by immunofluorescence analysis (Fig. 3A). No impact on parasite growth in culture conditions was observed (Fig. 3B). Next, we investigated whether the loss of FLAM8 could have impacted the total length of the flagellum based on the measurement of the signal obtained with the axonemal marker mAb25: no difference was observed among BSF lines (Fig. 3C). In addition, the absence of *FLAM8* did not affect parasite motility *in vitro*, neither in terms of speed nor in linearity (Fig. 3D-E, respectively). Compatible with our previous observations after RNAi silencing, these results show that pleomorphic trypanosomes tolerated the loss of *FLAM8 in vitro*.

**Figure 3.**
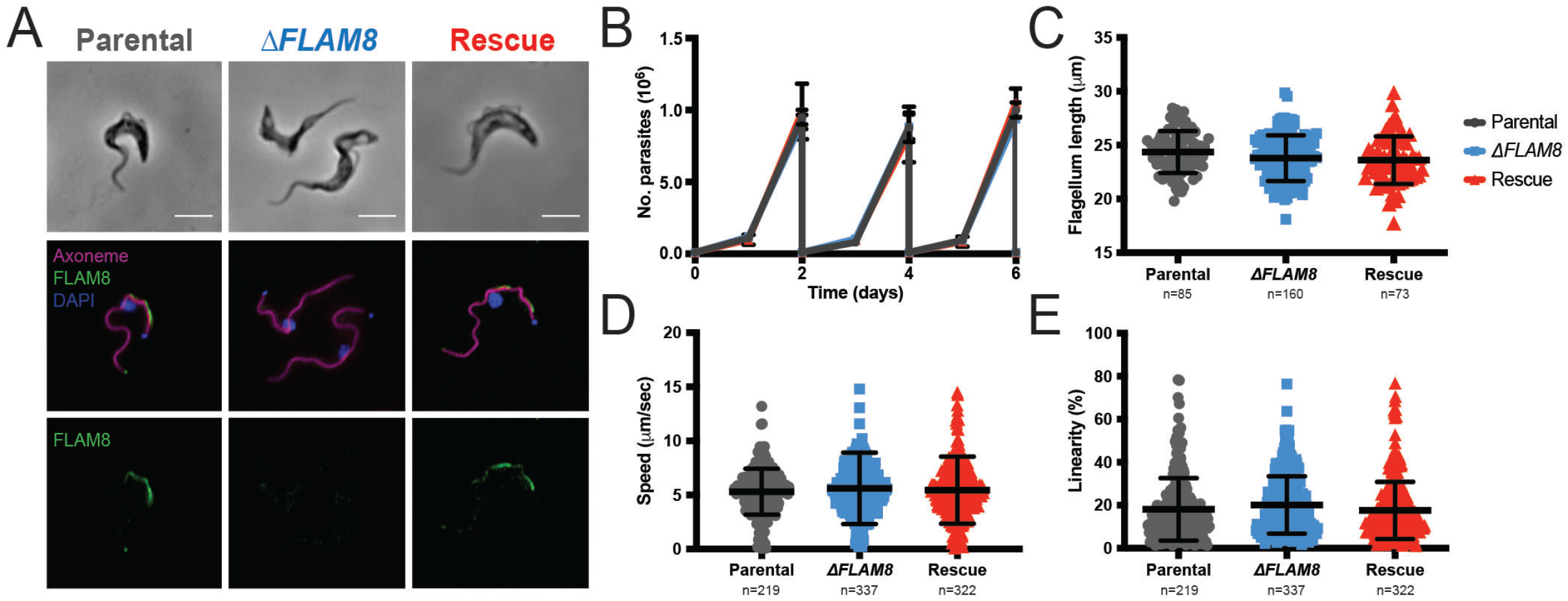
Characterization of the Δ*FLAM8* null mutants *in vitro*. **A)** Immunofluorescence pictures of parental, Δ*FLAM8* knockout and rescue pleomorphic BSF parasites labelled with the anti-FLAM8 (green) and mAb25 (axoneme in magenta) antibodies, DAPI staining for DNA content (blue). Scale bars show 5 μm. **B)** Growth curve of parental, Δ*FLAM8* and rescue pleomorphic BSF trypanosome cell lines over 6 consecutive days. **C)** Measurements of the flagellum length based on the axonemal marker mAb25 profiles in parental, Δ*FLAM8* and rescue parasites. No statistical differences were found. **D, E)** Motility tracking analysis showing the average speeds (D) and linearity (E) of BSF cell lines in matrix-dependent culture medium. No statistical differences were observed. The number of parasites considered for quantifications (N) is indicated below graphs (C), (D) and (E). Results represent the mean ± standard deviation (SD) of three independent experiments.

Then, functional investigations in the mammalian host were performed by infecting BALB/c mice either with the pleomorphic parental strain, three distinct Δ*FLAM8* knockout subclones or one Δ*FLAM8* strain bearing a rescue copy of *FLAM8* (Fig. 4). The infections were monitored daily during 4 weeks by quantifying the parasitemia and the bioluminescence signals emitted from whole animals (Fig. 4A). Null mutant parasites were able to establish an infection in the bloodstream as well as in the extravascular compartment. In the blood circulation, the absence of *FLAM8* did not prevent the *in vivo* differentiation of proliferative slender into transmissible stumpy parasites in the intravascular compartment (Fig. 4B), compatible with observations upon *in vitro* differentiation (Fig. S4, A-B). In addition, freshly-differentiated *FLAM8*-deprived stumpy parasites were able to further differentiate and maintain *in vitro* as procyclic trypanosomes (Fig. S4C). Within the vasculature, the overall amounts of parasites throughout the whole experimental infection were similar in all strains (Fig. 4C), yet significant differences between the three Δ*FLAM8* subclones as compared to parental-infected BALB/c mice were observed at days 5, 6 and 7, coinciding with the first peak of parasitemia (Fig. 4C). On the other hand, quantitative analyses of extravascular parasites showed a different scenario. Unlike in the intravascular compartment, significantly lower numbers of extravascular trypanosomes were observed between day 5 to 12, and from day 19 post-infection until the end of the experiment at day 27, evidencing an impaired extravascular colonization for Δ*FLAM8* parasites as compared to parental controls (Fig. 4D). To note, distribution profiles of all pleomorphic strains were different from those observed in mice infected with monomorphic parasites: larger amounts of trypanosomes were seen occupying the extravascular compartment reaching up to 5-8×10^9^ parasites, while the maximum amount found in the bloodstream never exceeded 7×10^7^ trypanosomes (Fig. S5).

**Figure 4.**
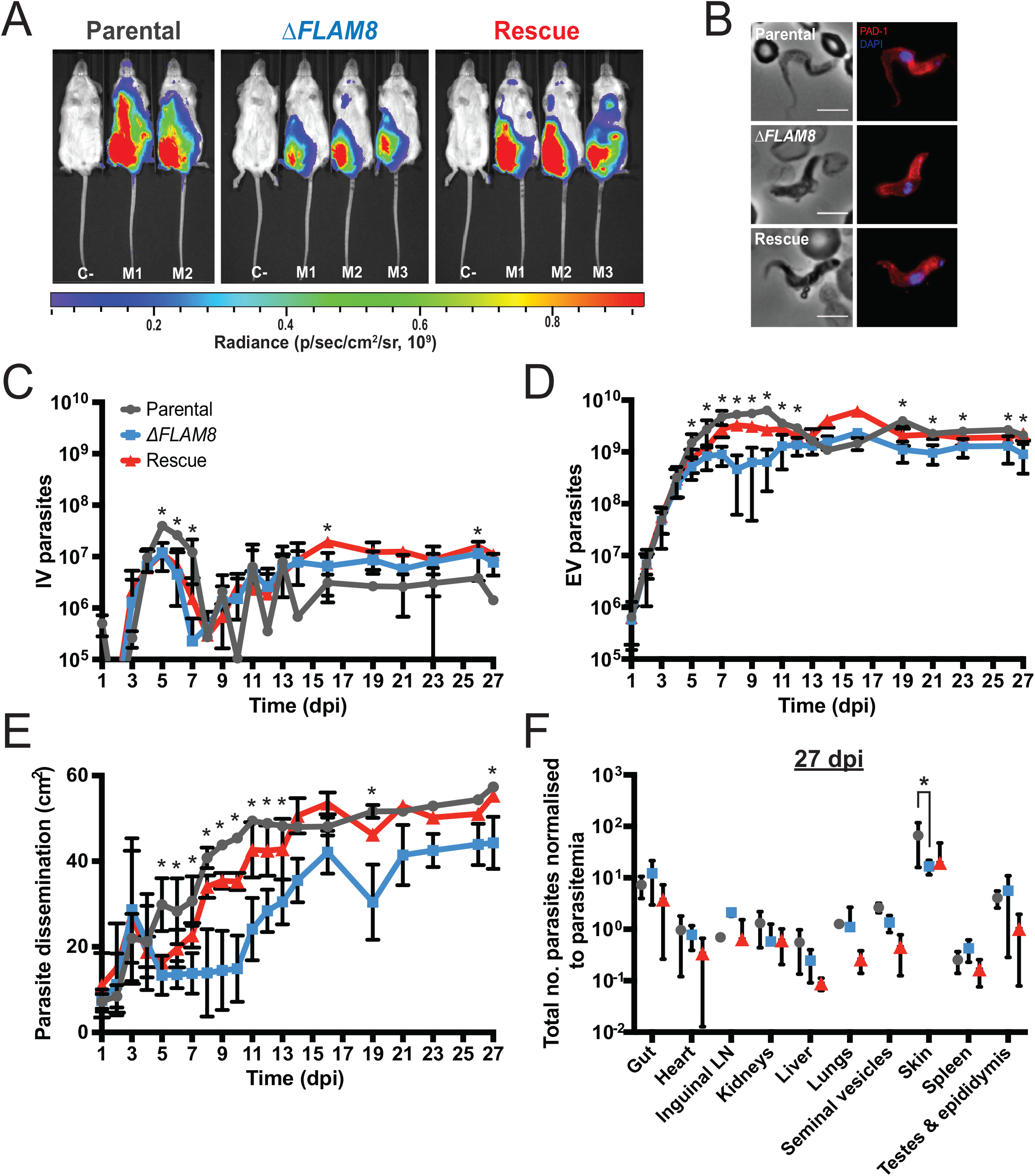
Functional investigations on the Δ*FLAM8* null mutants *in vivo* in the mammalian host. Groups of 3 BALB/c mice were injected intraperitoneally with either one parental, three Δ*FLAM8* null subclones or one rescue strains. One PBS-injected BALB/c animal was used as negative control. **A)** Normalized *in vivo* images of the bioluminescence radiance intensity (in photons / second / cm^2^ / steradian) emitted 8 days post-infection in BALB/c mice infected with parental, Δ*FLAM8* or rescue parasites (non-infected control mice C-were negative for bioluminescence). **B)** Representative immunofluorescence pictures of transmissible short stumpy parasites found in the blood of BALB/c mice infected with parental, Δ*FLAM8* and rescue parasites after the first peak of parasitemia. Anti-PAD1 antibody was used as a surface-exposed differentiation marker (red), DAPI staining for DNA content (blue). Scale bars represent 5 μm. **C)** Total number of parasites in the blood of infected mice (intravascular, IV) during the course of the infection (4 weeks). Statistically significant differences between the parental strain and Δ*FLAM8* subclones are indicated with *(p<0.0005). **D)** Total number of extravascular (EV) trypanosomes in the same mice. Statistically significant differences between the parental strain and Δ*FLAM8* subclones are indicated with *(p<0.0005). **E)** Dissemination of the parental, Δ*FLAM8* subclones and rescue parasite strains, measured over the entire animal body (in cm^2^) through the total surface of bioluminescent signal, during the entire infection course. Statistically significant differences between the parental strain and Δ*FLAM8* subclones are indicated with *(p<0.0005). Results represent means ± standard deviation (SD). **F)** Ratio between the total number of extravascular parasites quantified in individual dissected organs and the corresponding values of parasitemia at day 27 post infection in BALB/c mice infected with parental, Δ*FLAM8* null subclones or rescue pleomorphic BSF parasites. Only one statistically significant difference indicated with *(p<0.0005) was observed when comparing the skin of mice infected with Δ*FLAM8* and parental strains.

Furthermore, the subsequent quantification of the parasite spreading over the whole animal bodies showed that the depletion of *FLAM8* resulted in a partially yet significantly impaired dissemination of Δ*FLAM8* null mutant parasites in the extravascular compartment (days 5 to 13, 19 and 27 post-infection). This was mostly restored in trypanosomes bearing a rescue copy of *FLAM8* (Fig. 4E). Next, individual imaging of dissected organs was performed at the end of the experimental infection in order to estimate the amount of extravascular parasites in each organ. This was obtained by normalizing the total number of parasites within each organ (calculated from the bioluminescence) to the parasitemia level in the corresponding mouse at this last time point (Fig. 4F). At day 27, only the extravascular parasite colonization was significantly different between mice groups. Interestingly, the reduced distribution of *FLAM8*-deprived trypanosomes in the extravascular compartment was not detectable in most organs. Only the skin of Δ*FLAM8*-infected animals presented a statistically significant difference as compared to mice infected with the parental strain (Fig. 4F). In conclusion, these results indicate that FLAM8 is involved in the modulation of trypanosome distribution in the extravascular compartment of the mammalian host, especially in the skin.

## Discussion

The differential localization of FLAM8 from the very distal tip in tsetse midgut procyclic parasites to the entire flagellum length in the mammalian-infectious stages (Calvo-Alvarez *et al*., in revision) prompted us to speculate that FLAM8 could play a specific role in the mammalian host. Here, we present for the first time a connection of a flagellar protein with the efficiency of trypanosomes to distribute outside the host vasculature, especially in the skin. Quantitative analyses of experimental animal infections monitored by bioluminescence imaging showed that the absence of FLAM8 impaired parasite dissemination in the host extravascular compartment over the time of the infection, which was mostly recovered by the integration of a single rescue copy of the *FLAM8* gene in the endogenous locus.

### 1. FLAM8 and trypanosome transmission

Extravascular trypanosomes occupy the interstitial space of several organs, including the central nervous system, testes, adipose tissues and skin (Biteau et al., 2016; Caljon et al., 2016; Capewell et al., 2016; Carvalho et al., 2018; Goodwin, 1970; Trindade et al., 2016). The relevance of skin-dwelling trypanosomes in parasite transmission was demonstrated by xenodiagnosis experiments, early after the infective bite (Caljon et al., 2006), or later in the infection (Capewell et al., 2016), even in the absence of detectable parasitemia. More recently, the presence of extravascular trypanosomes was confirmed in the skin of confirmed and suspected cases of sleeping sickness (Camara et al., 2020). Here, the impaired spreading of *FLAM8*-null parasites over the extravascular compartment, especially in the skin, would mathematically reduce the probability for parasites to be ingested by tsetse flies.

In the bloodstream, the balance between proliferative slender parasites and tsetse-adapted stumpy forms responds to a quorum sensing mechanism involving the production of oligopeptides and their reception through a specific transporter (Rojas et al., 2018). Although stumpy proportions were not evaluated over the entire course of the infection, the absence of FLAM8 did not alter the ability of the parasites to differentiate into transmissible stumpy forms at the peak of parasitemia, showing that the protein is most probably not involved in this process. In addition, the ability of *FLAM8*-deprived stumpy BSF to differentiate into procyclics *in vitro* suggests that stumpy cells present in the blood and in the dermis could further develop upon ingestion by a tsetse fly.

### 2. On the possible cellular function(s) of FLAM8

Proliferative slender trypanosomes are highly mobile (Bargul et al., 2016) and this motility was proved to influence virulence *in vivo*. For instance, the lack of propulsive motility in flagellar dynein *LC1* knockout mutants resulted in the inability of trypanosomes to establish an infection in the bloodstream (Shimogawa et al., 2018). Here, *FLAM8* silencing in pleomorphic BSF trypanosomes did not alter either *in vitro* parasite growth or cell motility in matrix-dependent medium. However, quantitative analyses showed that *FLAM8*-null trypanosomes were less numerous in extravascular tissues as compared to parental controls. Assuming that parasite motility could be different in tissues and interstitial spaces of extravascular niches with biophysical properties distinct from those in the blood (Bargul et al., 2016; Sun et al., 2018), one cannot exclude that the motility of Δ*FLAM8* knockout parasites might be somehow altered in the extravascular compartment. Intravital imaging for motility analyses at the cell level would be needed in order to confirm this hypothesis.

Historically, the majority of studies on *T. brucei* virulence in experimental infections have considered the blood circulation almost as the sole host compartment parasitized by trypanosomes, whereas extravascular parasite niches and the underlying exchanges between both compartments have been underestimated for long. The fact that *FLAM8*-null trypanosomes were not able to disseminate properly over the extravascular compartment could somehow imply defects in the way parasites sense their microenvironment, resulting in an alteration of their extravascular tropism. For example, FLAM8 could be involved in the parasite extravasation process and / or in some subsequent host-parasite interactions promoting their maintenance and development once they have reached the extravascular compartment.

To our knowledge, FLAM8 is the first flagellar component affecting parasite dissemination in the extravascular host tissues *in vivo*. Further investigations on the FLAM8 interactions with other possible partners in the flagellum would help to unravel the roles of this fascinating and essential organelle, especially regarding the modulation of trypanosome distribution in their mammalian hosts, and its potential implications in parasite virulence and transmission.

## Experimental procedures

### Strains, culture and *in vitro* differentiation

The AnTat 1.1E Paris pleomorphic clone of *Trypanosoma brucei brucei* was derived from a strain originally isolated from a bushbuck in Uganda in 1966 (Le Ray, Barry, Easton, & Vickerman, 1977). The monomorphic *T. brucei* strain Lister 427 (Bohringer & Hecker, 1974) was also used. All bloodstream forms (BSF) were cultivated in HMI-11 medium supplemented with 10% (v/v) fetal bovine serum (FBS) (Hirumi & Hirumi, 1989) at 37°C in 5% CO_2_. For *in vitro* slender to stumpy BSF differentiation, we used 8-pCPT-2′-O-Me-5′-AMP, a nucleotide analog of 5’-AMP from BIOLOG Life Science Institute (Germany). Briefly, 2×10^6^ cultured pleomorphic AnTat 1.1E slender forms were incubated with 8-pCPT-2′-O-Me-5′-AMP (5 μM) for 48 h (Barquilla et al., 2012). For specific experiments, *in vitro* differentiation of BSF into PCF was performed by transferring freshly-differentiated short stumpy forms into SDM-79 medium supplemented with 10% (v/v) FBS, 6 mM cis-aconitate and 20 mM glycerol at 27°C (Czichos, Nonnengaesser, & Overath, 1986).

Monomorphic BSF “Single Marker” (SM) trypanosomes are derivatives of the Lister 427 strain, antigenic type MITat 1.2, clone 221a (Doyle, Hirumi, Hirumi, Lupton, & Cross, 1980), and express the T7 RNA polymerase and tetracycline repressor. *FLAM8^RNAi^* cells express complementary single-stranded RNA corresponding to a fragment of the *FLAM8* gene from two tetracycline-inducible T7 promoters facing each other in the pZJM vector (Wang, Morris, Drew, & Englund, 2000) integrated in the rDNA locus (Wirtz, Leal, Ochatt, & Cross, 1999). Addition of tetracycline (1 μg/mL) to the medium induces expression of sense and anti-sense RNA strands that can anneal to form double-stranded RNA (dsRNA) and trigger RNAi. For *in vivo* RNAi studies in mice, doxycycline hyclate (Sigma Aldrich) was added in sugared drinking water (0.2 g/L doxycycline hyclate combined with 50 g/L sucrose).

### Generation of FLAM8 RNAi mutants

For the generation of the *FLAM8^RNAi^* cell lines, a 380 bp (nucleotides 6665-7044) fragment of *FLAM8* (Tb927.2.5760), flanked by 5’ HindIII and 3’ XhoI restriction sites to facilitate subsequent cloning, was selected using the RNAit algorithm (http://trypanofan.bioc.cam.ac.uk/software/RNAit.html) in order to ensure that the targeted sequence was distinct from any other genes to avoid any cross-RNAi effects (Redmond, Vadivelu, & Field, 2003). This *FLAM8* DNA fragment was synthesized by GeneCust Europe (Dudelange, Luxembourg) and inserted into the HindIII-XhoI digested pZJM vector (Wang et al., 2000).

The pZJM-FLAM8 plasmid was linearized with NotI prior to transfection using nucleofector technology (Lonza, Italy) as described previously (Burkard, Fragoso, & Roditi, 2007). The cell line was further engineered for endogenous tagging of *FLAM8* with an mNeonGreen (mNG) at its C-terminal end by using the p3329 plasmid (Kelly et al., 2007), carrying a *FLAM8* gene fragment corresponding to *FLAM8* ORF nucleotides 8892-9391. Prior to nucleofection, NruI linearization of p3329-FLAM8-mNG plasmid was performed.

For *in vivo* experiments in mice, *FLAM8^RNAi^* parasites were finally modified by integrating a plasmid encoding for the chimeric triple reporter which combines the red-shifted firefly luciferase PpyREH9 and the tdTomato red fluorescent protein fused with a TY1 tag (Calvo-Alvarez et al., 2018). Transformants were selected with the appropriate antibiotic concentrations: phleomycin (1 μg/mL), blasticidin (5 μg/mL), G418 (2 μg/mL), and puromycin (0.1 μg/mL). Clonal populations were obtained by limiting dilution. Cell culture growth was monitored with an automatic Muse cell analyser (Merck Millipore, Paris).

### Generation of *FLAM8* KO mutants

For generating the *FLAM8* knockout and rescue cell lines, all insert templates were synthesized by GeneCust Europe (Dudelange, Luxembourg). For breaking the first allele, the 300 first nucleotides of the *FLAM8* gene flanking sequences were added on each side of a HYG resistance cassette (Fig. S2). For a complete disruption of the *FLAM8* locus, a second selectable marker (PAC) was flanked with the *FLAM8*-flanking sequence at 5’ and by 300 nucleotides of the *FLAM8* ORF (nucleotides 501-800) at 3’. For generating an add-back rescue cell line, due to the important size of the *FLAM8* ORF (9,228 nucleotides), the PAC selection marker was replaced by a BLE marker cassette flanked by the 300 first nucleotides of the *FLAM8* 5’ untranslated region (UTR) and by the nucleotides 1 to 500 of the *FLAM8* ORF for reinsertion into the endogenous locus of the knockout cell line. PCR amplifications of the DNA fragments bearing the *FLAM8* flanking sequences and the appropriate resistance markers were used for nucleofection and generation of all cell lines. The primers used are listed below: 5’-CATGACTTTACGTGTTTGGGCAC-3’ (FW, located 82 bp upstream the flanking 5’UTR sequence); 5’-CTTGCTTGTTTCTGTTTCGCAAC-3’ (RV, 130 bp downstream the flanking 3’UTR sequence, used to replace one WT allele by HYG resistance cassette); 5’-GCACACTAAAACTCATTGAAAGCC-3’ (RV, 926 bp downstream the ATG codon of *FLAM8*, used for second WT allele replacement by PAC cassette and rescue line generation). All knockout and rescue cell lines were further transfected to express the chimeric triple reporter protein PpyRE9H/TY1/tdTomato for multimodal *in vivo* imaging approaches as described elsewhere (Calvo-Alvarez et al., 2018). Selection-marker recovery was confirmed by screening individual clones in multi-well plates. Transformants were selected with the appropriate antibiotic concentrations: phleomycin (1 μg/mL), blasticidin (5 μg/mL), puromycin (0.1 μg/mL) and hygromycin (2.5 μg/mL). Clonal populations were obtained by limiting dilution and cell culture growth was monitored with an automatic Muse cell analyzer (Merck Millipore, Paris). Knockout and rescue cell lines were validated by whole-genome sequencing (BGI, Hong Kong). Briefly, genomic DNA from parental and mutant cell lines were sequenced by the HiSeq4000 sequence system (Illumina), generating about 10 million 100-bp reads and compared to that of the *T. brucei brucei* AnTat 1.1E Paris reference strain. In addition, some validation of the construct integrations in mutants were performed by PCR analysis according to standard protocols (Fig. S2 and Table 1).

**Table 1.**
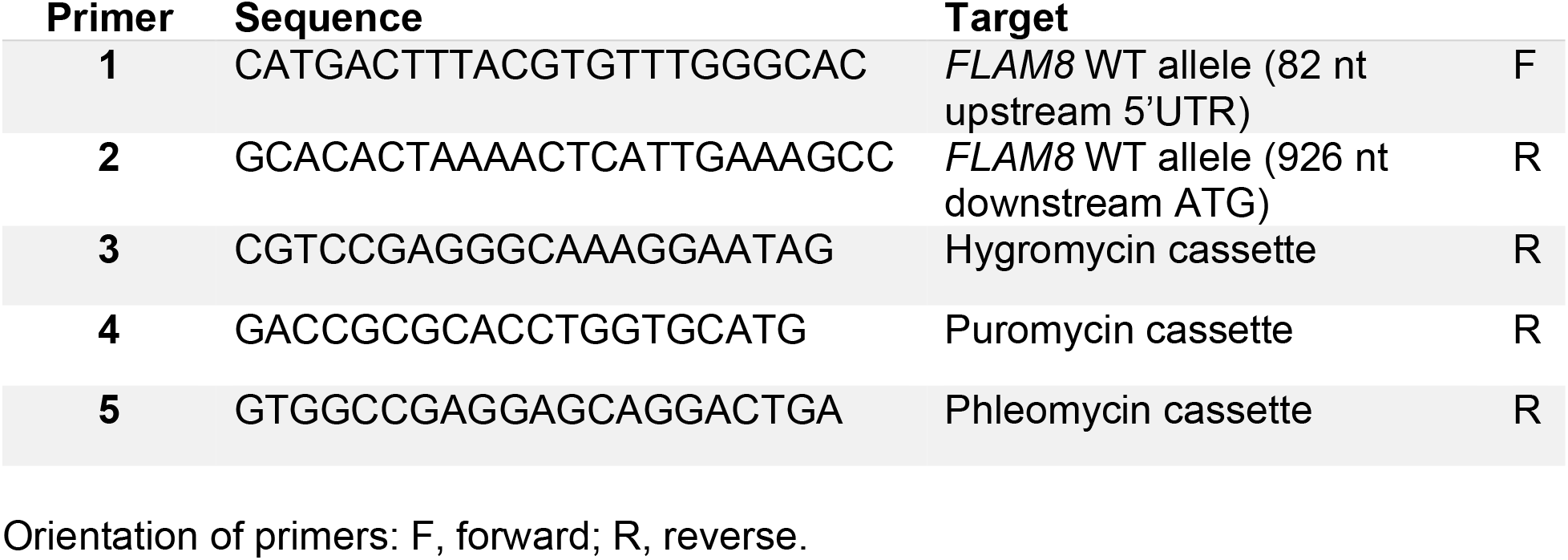
Oligonucleotides used for PCR validation of the Δ*FLAM8* knockout and rescue cell lines.

### Motility analyses

*In silico* 2D tracking was performed as previously described (Rotureau, Ooi, Huet, Perrot, & Bastin, 2014). For each BSF strain, 10 to 20 movies were recorded for 20 seconds (50 ms of exposure). Trypanosomes at 1×10^6^ parasites/mL were maintained in matrix-dependent HMI-11 medium containing 0,5% methylcellulose at 37°C and were observed under the 10x objective of an inverted DMI-4000B microscope (Leica) coupled to an ORCA-03G (Hamamatsu) or a PRIM95B (Photometrics) camera. Movies were converted with the MPEG Streamclip V.1.9b3 software (Squared 5) and analyzed with the medeaLAB CASA Tracking V.5.5 software (medea AV GmbH). Results were analyzed as mean ± SD of three independent experiments.

### *In vitro* bioluminescence quantification and analysis

To perform the parasite density / bioluminescence intensity assay, BSF parasites were counted, centrifuged and resuspended in fresh HMI-11 medium. Then, 100 μL of this suspension containing 10^6^ parasites were transferred into black clear-bottom 96-well plates and serial 2-fold dilutions were performed in triplicate adjusting the final volume to 200 μL with 300 μg/mL of beetle luciferin (Promega, France). Luciferase activity was quantified after 10 min of incubation with an IVIS Spectrum imager (PerkinElmer). Imaging data analysis was performed with the Living Image 4.3.1 software (PerkinElmer) by drawing regions of interest with constant size for well plate quantification. Total photon flux was calculated after removal of intensity values from WT parasites and / or parasite-free medium corresponding to the background noise. Results were analyzed as mean ± SD of three independent experiments.

### Mouse infection and ethical statements

Seven-week-old male BALB/c mice were purchased from Janvier Laboratory (sub-strain BALB/cAnNRj) and used as models for experimental infection and monitoring of the bioluminescence signal with the IVIS Spectrum imager (PerkinElmer). This study was conducted in strict accordance with the recommendations from the Guide for the Care and Use of Laboratory Animals of the European Union (European Directive 2010/63/UE) and the French Government. The protocol was approved by the “Comité d’éthique en expérimentation animale de l’Institut Pasteur” CETEA 89 (Permit number: 2012-0043 and 2016-0017) and undertaken in compliance with the Institut Pasteur Biosafety Committee (protocol CHSCT 12.131). BR is authorized to perform experiments on vertebrate animals (license #A-75-2035) and is responsible for all the experiments conducted personally or under his supervision. For *in vivo* infections, groups of four and three animals (*FLAM8* knockdown and knockout-infected mice, respectively) were injected intraperitoneally (IP) with 10^5^ slender BSF parasites, washed in TDB (Trypanosome Dilution Buffer: 5 mM KCl, 80 mM NaCl, 1 mM MgSO_4_*7H_2_O, 20 mM Na_2_HPO_4_, 2 mM NaH_2_PO_4_, 20 mM glucose) and resuspended in 100 μl of PBS prior animal inoculation.

### *In vivo* bioluminescence imaging, analysis and parasitemia quantifications

Infection with bioluminescent parasites was monitored daily by detecting the bioluminescence signal in whole animals with the IVIS Spectrum imager (PerkinElmer). The equipment consists of a cooled charge-coupled camera mounted on a light-tight chamber with a nose cone delivery device to keep the mice anaesthetized during image acquisition with 1.5-2% isoflurane. A heated stage is comprised within the IVIS Spectrum imager in order to maintain optimum body temperature. D-luciferin potassium salt (Promega) stock solution was prepared in sterile PBS at 33.33 mg/mL, and stored in a −20°C freezer. To produce bioluminescence, mice were inoculated by the intraperitoneal route with 150 μL of D-luciferin stock solution (250 mg/Kg body weight). After 10 minutes of incubation to allow substrate dissemination, all mice were anaesthetized in an oxygen-rich induction chamber with 1.5-2% isoflurane, and images were acquired by using automatic exposure (30 seconds to 5 minutes) depending on signal intensity.

Images were analyzed with the Living Image software version 4.3.1 (PerkinElmer). Data were expressed in total photons/second (p/s) corresponding to the total flux of bioluminescent signal according to the selected area (regions of interest with constant size covering the total body of the mouse). The background noise was removed by subtracting the bioluminescent signal of the control mouse from the infected ones for each acquisition. For parasite dissemination analyses, a minimum value of photons/second (p/s) was set for all animals in every time point in order to quantify the exact dissemination area (in cm^2^) over the whole animal body. Parasitemia was determined daily following tail bleeds and assayed by automated fluorescent cell counting with a Muse cytometer (Merck-Millipore, detection limit at 10^2^ parasites/mL) according to the manufacturer’s recommendations. The quantification of the total intravascular parasite population was assessed by calculating the blood volume of all animals according to their body weight and referring to daily parasitemia. To quantify the number of extravascular parasites, a region of interest with a constant size was used to obtain the total number of parasites within the entire animal bodies (total bioluminescence). Subsequently, the total number of parasites present in the vascular system was subtracted, resulting in estimating the total parasite population colonizing the extravascular compartments at a given time point.

### Immunofluorescence analysis (IFA)

Cultured parasites were washed twice in TDB and spread directly onto poly-L-lysine coated slides. For methanol fixation, slides were air-dried for 10 min, fixed in methanol at −20°C for 5 min and rehydrated for 20 min in PBS. For immunodetection, slides were incubated for 1 h at 37°C with the appropriate dilution of the first antibody in 0.1% BSA in PBS. After 3 consecutive 5 min washes in PBS, species and subclass-specific secondary antibodies coupled to the appropriate fluorochrome (Alexa 488, Cy3, Cy5 Jackson ImmunoResearch) were diluted 1/400 in PBS containing 0.1% BSA and were applied for 1 h at 37°C. After washing in PBS as indicated above, slides were finally stained with 4’,6-diamidino-2-phenylindole (DAPI, 1 μg/mL) for visualization of kinetoplast and nuclear DNA content, and mounted under cover slips with ProLong antifade reagent (Invitrogen), as previously described (Rotureau et al., 2011). Slides were observed under an epifluorescence DMI4000 microscope (Leica) with a 100x objective (NA 1.4), an EL6000 (Leica) as light excitation source and controlled by the Micro-Manager V1.4.22 software (NIH), and images were acquired using an ORCA-03G (Hamamatsu) or a PRIM95B (Photometrics) camera. Images were analyzed with ImageJ V1.8.0 (NIH). The monoclonal antibody mAb25 (anti-mouse IgG2a, 1:10) was used as a flagellum marker as it specifically recognizes the axoneme protein *Tb*SAXO1 (Dacheux et al., 2012). FLAM8 was detected using: i) a specific rabbit serum (1:500) kindly provided by Paul McKean (University of Lancaster, UK), or ii) a monoclonal anti-mNeonGreen antibody (anti-mouse IgG2c, 1:100, ChromoTek). Stumpy BSF were identified at the molecular level with a rabbit polyclonal anti-PAD1 antibody (kindly provided by Keith Matthews, University of Edinburgh; dilution 1:300) (Dean, Marchetti, Kirk, & Matthews, 2009). In the case of RNAi knockdown experiments, IFA signals were normalized using the signal obtained in non-induced controls as a reference.

### Measurements, normalization and statistical analyses

Standardization of fluorescent signals was carried out by parallel setting of raw integrated density signals in all the images to be compared in ImageJ V1.8.0 (NIH). For clarity purposes, the brightness and contrast of several pictures were adjusted after their analysis in accordance to editorial policies. Statistical analyses and plots were performed with XLSTAT 2019.2.01 (Addinsoft) in Excel 2016 (Microsoft) or Prism V8.2.1 (GraphPad). Statistical analyses include linear regression analyses for bioluminescence / fluorescence intensity vs. parasite density, two-sided ANOVA / ANCOVA tests for growth curves, with Tukey ad-hoc post-tests for inter-group comparison, all at 95% confidence.

## Acknowledgements

We thank M. Bonhivers, D. Robinson, P. McKean, K. Matthews and K. Gull for providing various plasmids and antibodies. We acknowledge the IP BioImaging Plateforme for providing access to their equipment. We are grateful to P. Bastin for his strong scientific and human support. We warmly thank P. Bastin, M. Boshart and S. Bonnefoy for their critical reading of the manuscript.

## Author contributions

ECA, CT and AC performed the experiments. ECA and BR designed the study, analyzed the data and wrote the manuscript.

