## Supplementary material for "FLAgellum Member 8 modulates extravascular trypanosome distribution in the mammalian host"

**Supplementary figures and legends**

**Figure S1. Validation of the triple-reporter efficiency in monomorphic *FLAM8*$^{RNAi}$ parasite lines.** Linear correlation between the number of parasites and the bioluminescence (in p/s) emitted by monomorphic *FLAM8*$^{RNAi}$ *FLAM8::mNG* 427 BSF overexpressing the triple reporter chimeric protein (Calvo-Alvarez et al., 2018) acquired by the IVIS Spectrum imager. Parasites without the RNAi plasmid (control), non-treated with tetracycline (non-induced) and treated with tetracycline (induced) are shown. Representative bioluminescent image of serial 1/2 dilutions performed in a 96-well plate (in photons / second / cm$^2$ / steradian). RNAi induction was triggered by the addition of 1 µg tetracycline and / or doxycycline for 72 h. Results represent the mean ± standard deviation (SD) of three independent experiments.

**Figure S2. Molecular validation of the Δ*FLAM8* null mutant cell lines. A)** Whole-genome sequencing results showing *FLAM8* wild-type allele (WT, upper panel), the loss of part of the 5' *FLAM8* ORF in Δ*FLAM8* knockout trypanosomes (middle panel) and the restoration of the full *FLAM8* gene in rescue parasites (bottom panel), relative to the number of reads per 100-nt read length. The presence of the correct antibiotic cassettes is shown for knockout and rescue parasites (right middle and bottom panels). Δ*FLAM8* knockout and rescue parasites also bear a construct for expression of a triple reporter (TR) as assessed by the detection of *bsd* reads. **B)** Schemes showing the structure of the *FLAM8* locus in wild-type parasites (upper scheme) and the integration plan of the different cassettes (lower *HYG*-, *PAC*- and *BLE*-containing schemes). PCR confirmation of the successful integrations of all reporter cassettes (right panel). Primer pairs used for PCRs are indicated at the bottom of each line and correspond to those drawn on the schemes. BSD: blasticidin; HYG: hygromycin; PAC: puromycin; BLE: phleomycin.

**Figure S3. Validation of the triple-reporter efficiency in pleomorphic Δ*FLAM8* mutant cell lines.** Linear correlation between the number of parasites and the bioluminescence (in p/s) emitted by pleomorphic parental, Δ*FLAM8* knockout subclones and rescue parasites overexpressing the triple reporter chimeric protein (Calvo-Alvarez et al., 2018) acquired by the IVIS Spectrum imager. Representative bioluminescent image of serial 1/2 dilutions performed in a 96-well plate (in photons / second / $cm^2$ / steradian). Results represent the mean ± standard deviation (SD) of three independent experiments.

**Figure S4. The absence of *FLAM8* does not impact the parasite differentiation into transmissible stumpy and insect procyclic forms *in vitro*. A)** Representative immunofluorescence pictures of methanol-fixed parental (left panel), Δ*FLAM8* null subclones (middle panels) and rescue (right panel) stumpy trypanosomes, upon *in vitro* treatment with a nucleotide 5'-AMP analog. Stumpy BSF were stained with anti-PAD1 antibody (red) and DAPI for DNA content (blue). Scale bars represent 5 µm. **B)** Quantification of the proportion of stumpy trypanosomes in all pleomorphic cell lines after *in vitro* differentiation in three independent experiments. No significant statistical differences were found. The number of parasites considered for quantification (N) is indicated above the graph. **C)** Representative pictures of methanol-fixed procyclic parasites after *in vitro* differentiation from parental (left panel), Δ*FLAM8* null subclones (middle panels) and rescue (right panel) trypanosomes. Parasites were stained with the anti-axonemal mAb25 (magenta) and anti-FLAM8 (green) antibodies, DAPI staining for DNA content (blue). Scale bars represent 5 µm.

**Figure S5. Comparison of intra- and extravascular trypanosome populations during the experimental infection with the Δ*FLAM8* null mutants. A-E)** Total numbers of parasites present in the bloodstream (IV, continuous line) and in the extravascular tissues (EV, dotted line) of BALB/c mice infected with pleomorphic parental (A), rescue (B) and three Δ*FLAM8* null subclones (C, D, E) during the whole course of the infection (27 days). Statistically significant differences were found in all parasite strains during the entire infection period. The presence of parasites by bioluminescence imaging was evident at day 1 post-infection, while blood parasites were first detected at day 3, in agreement with previous observations (Capewell et al., 2016).

### Figure S1

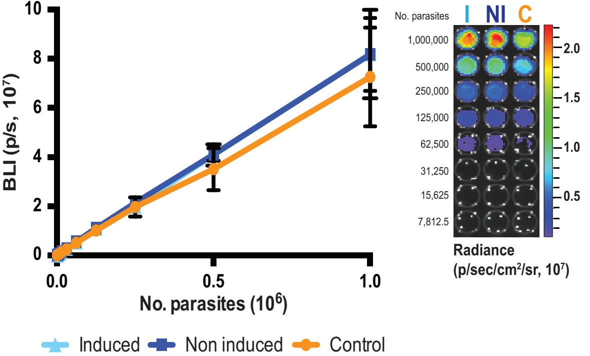
BLI (p/s, 10^7)
10
8
6
4
2
0
0.0
0.5
1.0
No. parasites (10^6)
Induced
Non induced
Control
No. parasites
I
NI
C
1,000,000
500,000
250,000
125,000
62,500
31,250
15,625
7,812.5
2.0
1.5
1.0
0.5
Radiance
(p/sec/cm²/sr, 10^7)

### Figure S2

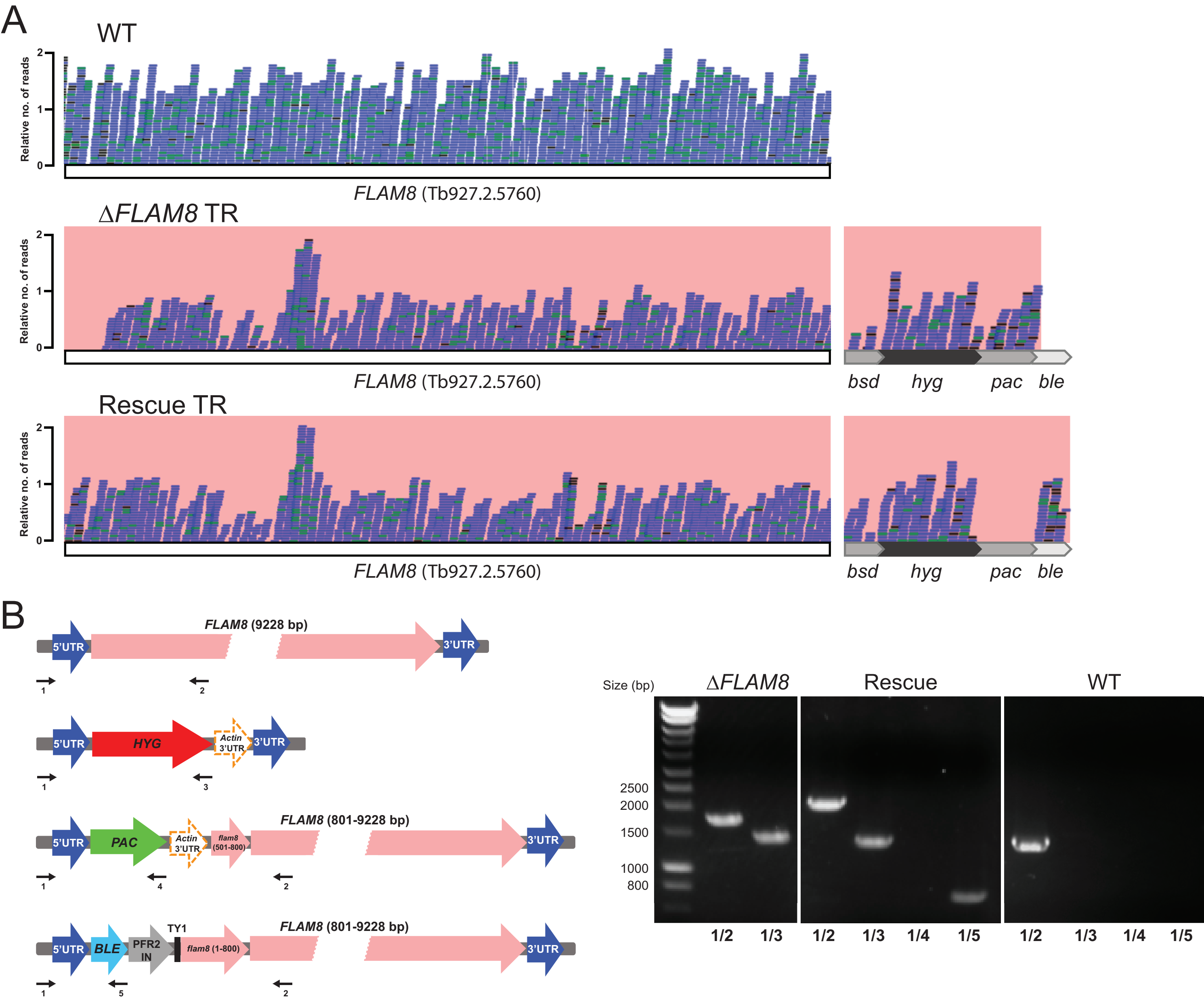
A
WT
Relative no. of reads
2
1
0
FLAM8 (Tb927.2.5760)
ΔFLAM8 TR
Relative no. of reads
2
1
0
FLAM8 (Tb927.2.5760)
bsd
hyg
pac
ble
Rescue TR
Relative no. of reads
2
1
0
FLAM8 (Tb927.2.5760)
bsd
hyg
pac
ble
B
FLAM8 (9228 bp)
5'UTR
3'UTR
1
2
5'UTR
HYG
Actin 3'UTR
3'UTR
1
3
FLAM8 (801-9228 bp)
5'UTR
PAC
Actin 3'UTR
flam8 (501-800)
3'UTR
1
4
2
TY1
FLAM8 (801-9228 bp)
5'UTR
BLE
PFR2 IN
flam8 (1-800)
3'UTR
1
5
2
Size (bp)
ΔFLAM8
Rescue
WT
2500
2000
1500
1000
800
1/2
1/3
1/2
1/3
1/4
1/5
1/2
1/3
1/4
1/5

### Figure S3

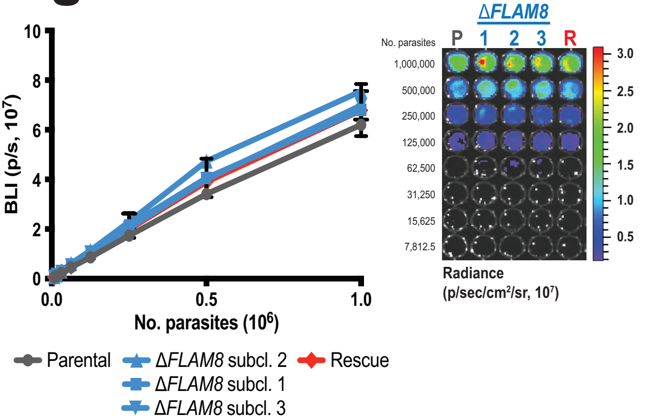

BLI (p/s, 10^7)
10
8
6
4
2
0
0.0
0.5
1.0
No. parasites (10^6)
Parental
ΔFLAM8 subcl. 2
ΔFLAM8 subcl. 1
ΔFLAM8 subcl. 3
Rescue
ΔFLAM8
P 1 2 3 R
No. parasites
1,000,000
500,000
250,000
125,000
62,500
31,250
15,625
7,812.5
3.0
2.5
2.0
1.5
1.0
0.5
Radiance
(p/sec/cm²/sr, 10⁷)

### Figure S4

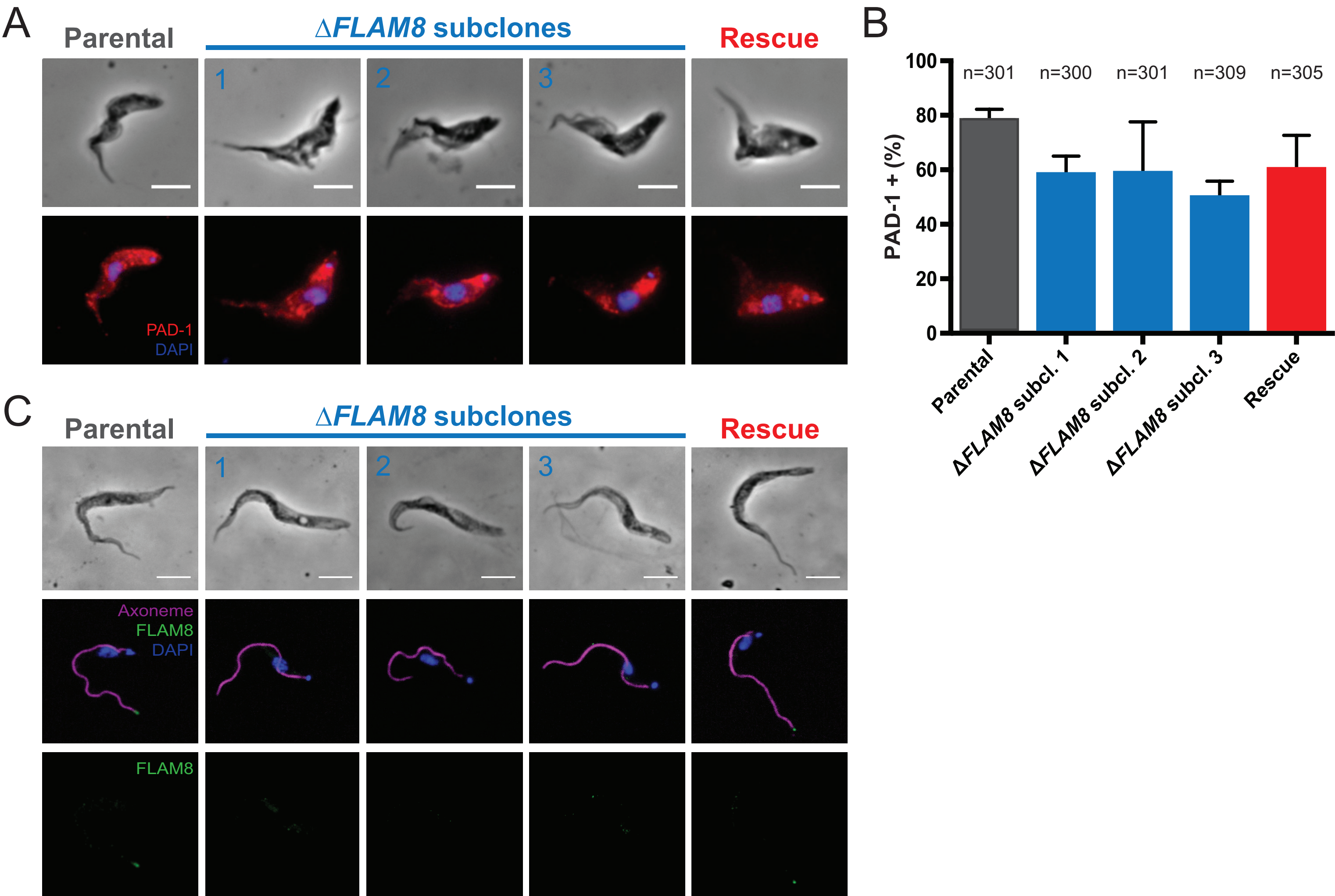

A
Parental
ΔFLAM8 subclones
Rescue
1
2
3
PAD-1
DAPI
B
100
80
60
40
20
0
PAD-1 + (%)
n=301
n=300
n=301
n=309
n=305
Parental
ΔFLAM8 subcl. 1
ΔFLAM8 subcl. 2
ΔFLAM8 subcl. 3
Rescue
C
Parental
ΔFLAM8 subclones
Rescue
1
2
3
Axoneme
FLAM8
DAPI
FLAM8

### Figure S5

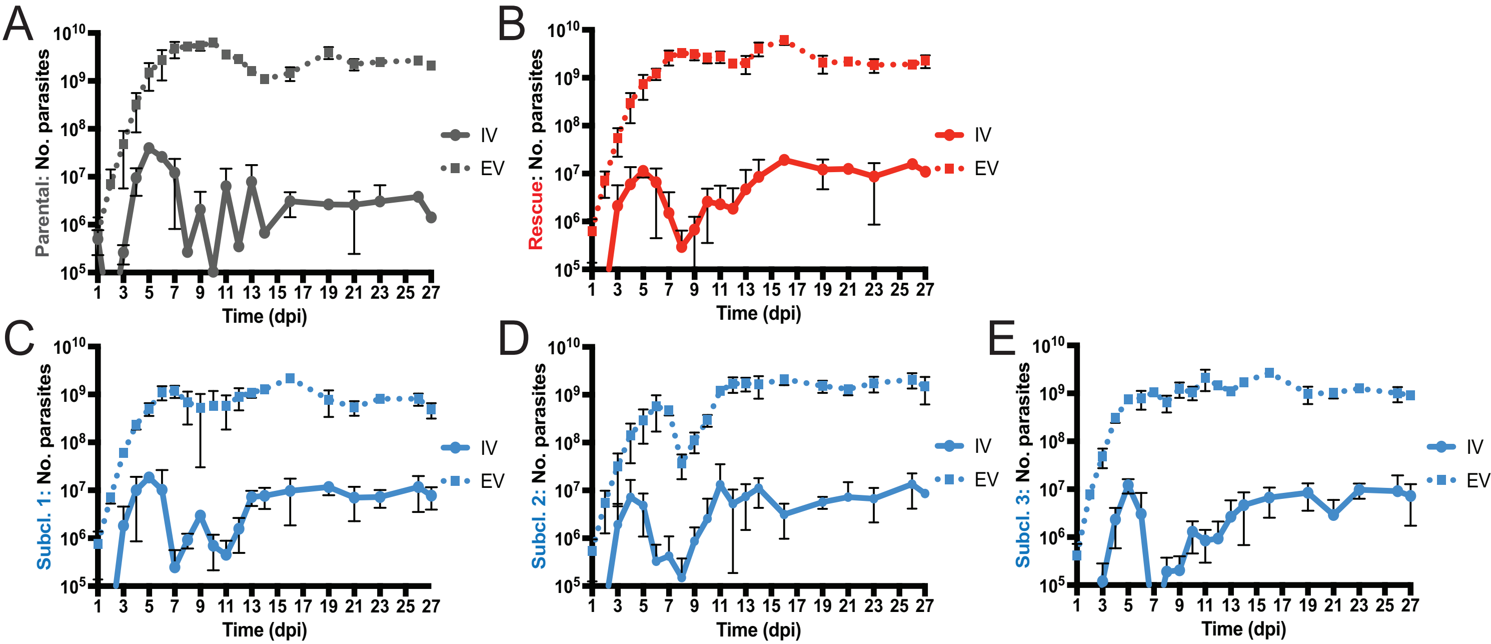
A
B
C
D
E
Parental: No. parasites
Rescue: No. parasites
Subcl. 1: No. parasites
Subcl. 2: No. parasites
Subcl. 3: No. parasites
$10^{10}$
$10^{9}$
$10^{8}$
$10^{7}$
$10^{6}$
$10^{5}$
1 3 5 7 9 11 13 15 17 19 21 23 25 27
Time (dpi)
IV
EV
